# The Spliceosome Scaffold Component BCAS2 Promotes Pancreatic Cancer Progression through Weak 5′ Splice Site–Dependent Splicing Regulation

**DOI:** 10.64898/2026.10.06.757127

**Authors:** Ledong Wan, Mei Wang, Yuma Ishigami, Alexander J. Kral, Kuan-Ting Lin, Joseph R. Merrill, Scott K. Lyons, Adrian R. Krainer

**Affiliations:** Department of Pharmacological Sciences, State University of New York at Stony Brook, Stony Brook, NY 11794, USA; Stony Brook Cancer Center, Renaissance School of Medicine at Stony Brook University, Stony Brook, NY 11794, USA; Cold Spring Harbor Laboratory, Cold Spring Harbor, NY 11724, USA

## Abstract

Pancreatic ductal adenocarcinoma (PDAC) remains one of the deadliest malignancies. Dysregulated RNA splicing has emerged as a critical mechanism contributing to PDAC progression. However, whether structural components of the spliceosome regulate oncogenic alternative splicing was not known. Here, we identify the nineteen complex (NTC) component BCAS2/SPF27 as a critical regulator of PDAC progression. Elevated expression of BCAS2 in PDAC is associated with poor patient outcomes. Transcriptomic analyses revealed that BCAS2 specifically regulates alternative exons with weak splice sites. In particular, BCAS2 promotes *SOS1* exon 21 inclusion and consequently RAC-signaling activation. Furthermore, splice-switching antisense oligonucleotide (ASO)-mediated skipping of *SOS1* exon 21 suppressed PDAC proliferation and migration. Our findings uncover a role of BCAS2 in the regulation of weak splice-site usage and identify the BCAS2–SOS1–RAC signaling axis as a potential therapeutic vulnerability in pancreatic cancer.

## INTRODUCTION

Pancreatic ductal adenocarcinoma (PDAC) remains one of the most lethal malignancies, with a five-year survival rate below 13%.^1^ Despite recent advances in targeted therapies, most patients are diagnosed with advanced-stage PDAC and respond poorly to current treatments, because of extensive molecular heterogeneity and the rapid emergence of therapeutic resistance.^2–4^ In addition to somatic mutations, dysregulated alternative RNA splicing markedly contributes to PDAC progression.^5–7^ Aberrant alternative splicing generates oncogenic isoforms, alters signaling pathways, and reprograms the tumor microenvironment to support malignancy.^8, 9^

The spliceosome, a large ribonucleoprotein complex responsible for pre-mRNA splicing, consists of small nuclear ribonucleoproteins (snRNPs) and numerous associated proteins that coordinate splice-site recognition, spliceosome remodeling, and catalysis.^10, 11^ Increasing evidence suggests that cancer cells exploit alterations in the splicing machinery to promote malignant progression. Recurrent mutations in spliceosomal components and splicing regulators, including SF3B1^12–14^, U2AF1^15, 16^, and SRSF2^17, 18^, as well as dysregulated expression of splicing factors such as SRSF1^5, 19, 20^, promote oncogenic alternative splicing across multiple cancers. However, whether non-mutated structural components of the spliceosome actively regulate oncogenic splicing remains largely unexplored.

During spliceosome activation, the nineteen complex (NTC) is recruited after U4 snRNP dissociation and functions as a scaffold to stabilize interactions among U5, U6, and U2 snRNPs, promoting the conformational rearrangements required for catalytic activation and the two transesterification reactions of splicing.^21, 22^ BCAS2 (breast-cancer amplified sequence at Chromosome 1p13.2), also known as Splicing Factor 27 (SPF27) and DAM1 (DNA amplified in mammary carcinoma), encodes a core component of the NTC complex, within which it associates with PRPF19 and CDC5L.^23–26^ Genetic studies highlight its essential and conserved role in RNA processing, as loss of *dBCAS2* in flies causes defective RNA splicing, abnormal wing development, and lethality that can be partially rescued by human BCAS2.^27^ Consistently, homozygous deletion of *Bcas2* is embryonically lethal in mice.^28^ BCAS2 also regulates specific alternative splicing, including *Delta* pre-mRNA splicing in the Notch signaling pathway and splicing events required for mouse spermatogenesis.^28, 29^

Beyond its canonical role in spliceosome regulation, BCAS2 has been implicated in cancer progression. BCAS2 was originally identified as an amplified gene in breast cancer and is overexpressed in multiple malignancies.^30, 31^ Previous studies demonstrated that BCAS2 cooperates with estrogen receptor alpha (ERα) to promote breast-cancer progression, enhances androgen-receptor expression in prostate cancer, and interacts with Nibrin/NBS1 to facilitate DNA double-strand break repair.^32–34^ However, these studies have largely focused on non-splicing functions of BCAS2. Given its fundamental role as an NTC scaffold component in spliceosome activation, whether cancer cells exploit BCAS2 to drive oncogenic alternative splicing programs remained unknown.

In this study, we identify BCAS2 as a critical regulator of alternative RNA splicing in pancreatic cancer. We demonstrate that BCAS2 expression is upregulated in human PDAC tumors and correlates with poor patient outcomes. BCAS2 promotes PDAC cell proliferation and migration *in vitro*, and tumor growth and metastasis *in vivo*. Transcriptomic and mechanistic analyses demonstrate that BCAS2 preferentially regulates alternative exons with weak splice sites, thereby controlling a subset of cancer-associated splicing events that converge on RHO signaling. Our findings identify a previously unrecognized role of BCAS2 in PDAC and highlight splicing modulation as a potential therapeutic opportunity.

## RESULTS

### Increased expression of BCAS2 is associated with adverse PDAC clinical outcomes

We first examined whether NTC components are dysregulated in PDAC. We found that *BCAS2*, *PRPF19*, and *CDC5L* are all upregulated in PDAC tumors, compared with normal pancreatic tissues (Fig. 1A and S1A). However, only high *BCAS2* expression is significantly associated with poor patient outcomes (Fig. 1B and S1A). Notably, *BCAS2* expression is relatively low in normal pancreatic tissues, compared with other normal tissues (Fig. S1B). Among different cancer types, pancreatic cancer, diffuse large B-cell lymphoma, and thymoma exhibit significantly upregulated *BCAS2* expression (Fig. S1C). Immunohistochemical (IHC) staining confirmed increased BCAS2 protein levels in human PDAC tissues and the KPC (*LSL-Kras^G12D/+^*; *LSL-Trp53^R172H/+^*; *Pdx1-Cre*) mouse model^35^ (Fig. 1C and 1D).

**Figure 1.**
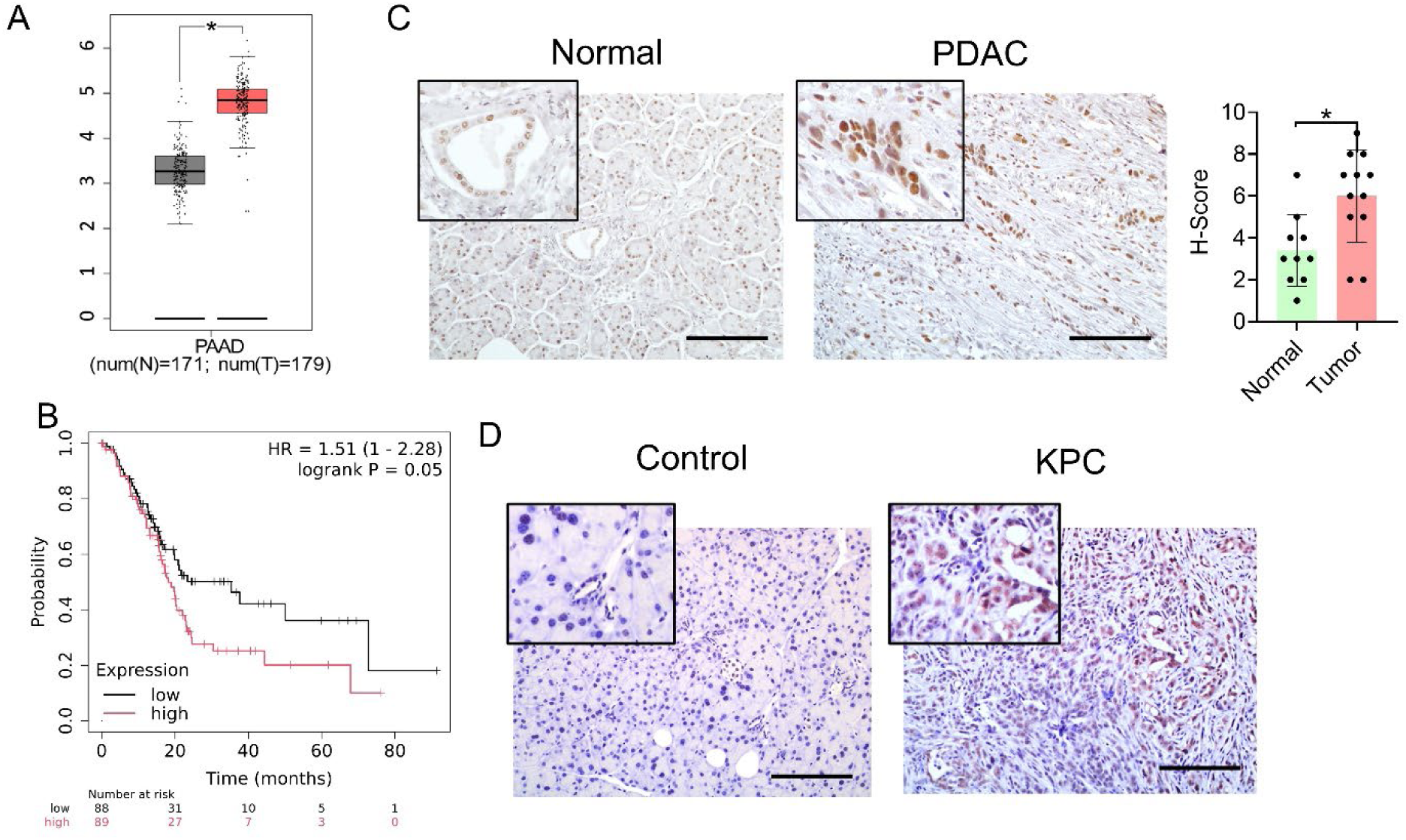
Elevated BCAS2 expression is associated with poor prognosis in PDAC. (A) *BCAS2* expression in normal pancreatic samples and pancreatic cancer from TCGA dataset. (B) Kaplan–Meier survival analysis of PDAC patients with different *BCAS2* expression. (C) IHC staining (left) and intensity scores (right) of BCAS2 in human normal pancreatic tissue (n = 10) and PDAC tumors (n = 12). Scale bars, 100 μm. Unpaired t test, *p <0.05. Error bars represent mean ± SD. (D) IHC staining of BCAS2 in normal mouse pancreatic tissue and KPC PDAC tumors. n = 5. Scale bars, 100 μm.

### BCAS2 promotes PDAC progression *in vitro*

We next analyzed BCAS2 protein levels in a panel of PDAC cell lines, and found that MiaPaCa- 2 and SUIT-2 cells exhibited high expression, whereas PANC-1 and AsPC-1 cells showed lower expression (Fig. 2A). We designed short hairpin RNAs (shRNAs) to knock down *BCAS2* expression in SUIT-2 and MiaPaCa-2 cells (Fig. 2B). *BCAS2* knockdown markedly suppressed anchorage-independent growth and cell migration (Fig. 2C and 2D). Conversely, overexpression of BCAS2 in PANC-1 cells promoted colony formation and migration (Fig. 2E and 2F).

**Figure 2.**
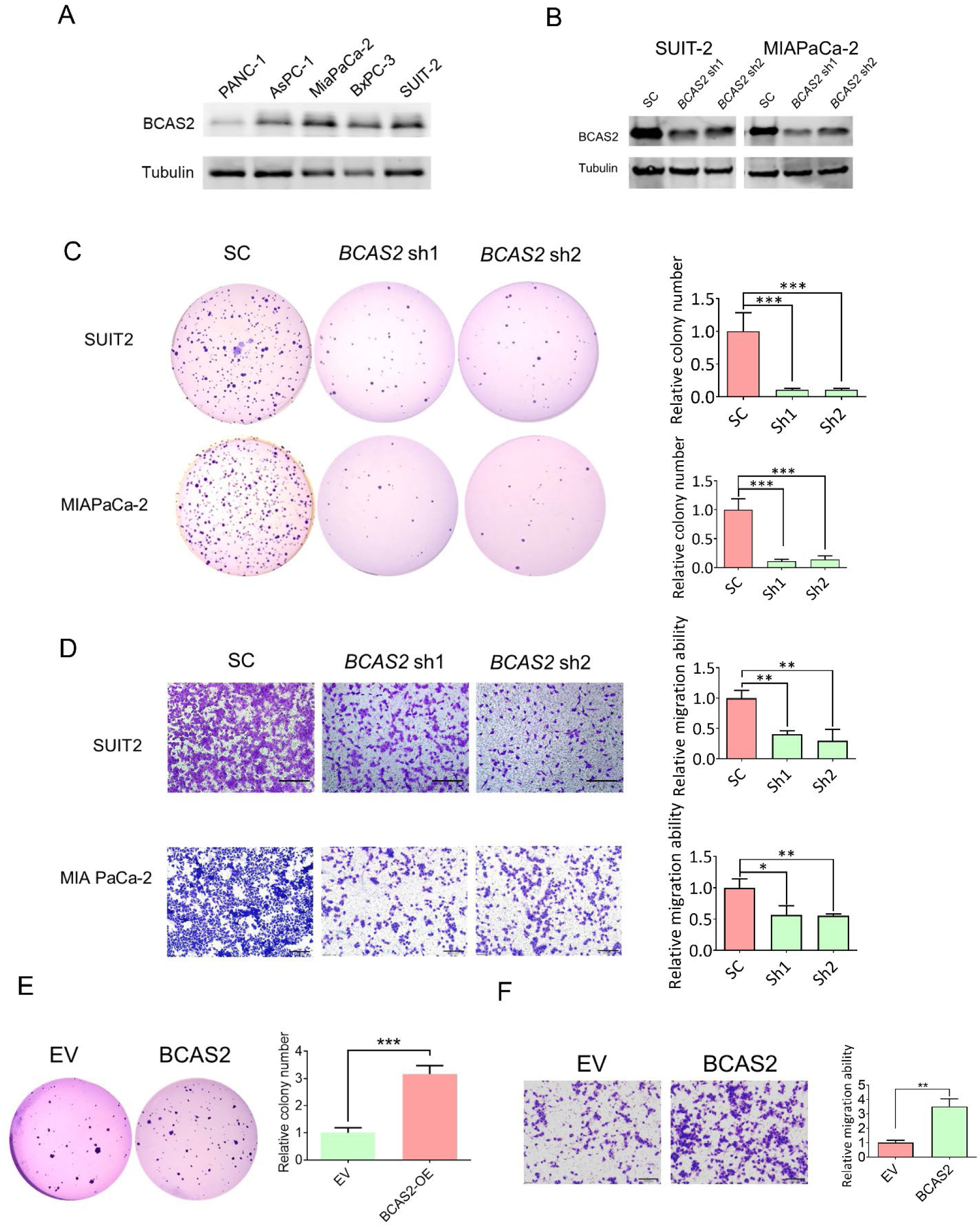
BCAS2 promotes PDAC tumor progression *in vitro*. (A) Western blotting of BCAS2 in various PDAC cell lines. (B) Western blotting of BCAS2 in MiaPaCa-2 and SUIT-2 cells with *BCAS2* knockdown. (C) Soft-agar colony-formation assay (left) and quantification (right) of MiaPaCa-2 and SUIT-2 cells with *BCAS2* knockdown. One-way ANOVA with Tukey multiple comparisons test. ***p < 0.001; n = 3. Error bars represent mean ± SD. (D) Transwell assay (left) and quantification (right) of the cell-migration properties of MiaPaCa-2 and SUIT-2 cells with *BCAS2* knockdown. One-way ANOVA with Tukey multiple comparisons test. *p < 0.05, **p < 0.01; n = 3. Error bars represent mean ± SD. Scale bars, 200 μm. (E) Soft-agar colony-formation assay (left) and quantification (right) of PANC-1 cells with *BCAS2* overexpression. One-way ANOVA with Tukey multiple comparisons test. ***p < 0.001; n = 3. Error bars represent mean ± SD. (F) Transwell assay (left) and quantification (right) of the cell-migration properties of PANC-1 cells with *BCAS2* overexpression. One-way ANOVA with Tukey multiple comparisons test. **p < 0.01; n = 3. Error bars represent mean ± SD. Scale bars, 200 μm.

### BCAS2 promotes PDAC tumor growth and metastasis *in vivo*

To evaluate the role of BCAS2 in PDAC tumor growth *in vivo*, we orthotopically transplanted MiaPaCa-2 cells stably expressing luciferase with either scramble control or *BCAS2* shRNAs into the pancreas of immunodeficient mice. Bioluminescence imaging revealed that *BCAS2* knockdown significantly suppressed tumor growth (Fig. 3A). At the endpoint, *BCAS2*-knockdown tumors exhibited significantly reduced size and weight (Fig. 3B). Additionally, metastatic lesions were detected in the lung and liver tissues of control mice, whereas no detectable metastatic signals were observed in *BCAS2*-knockdown mice (Fig. 3C). Accordingly, circulating tumor cells (CTCs) were markedly reduced in *BCAS2*-knockdown mice (Fig. 3D).

**Figure 3.**
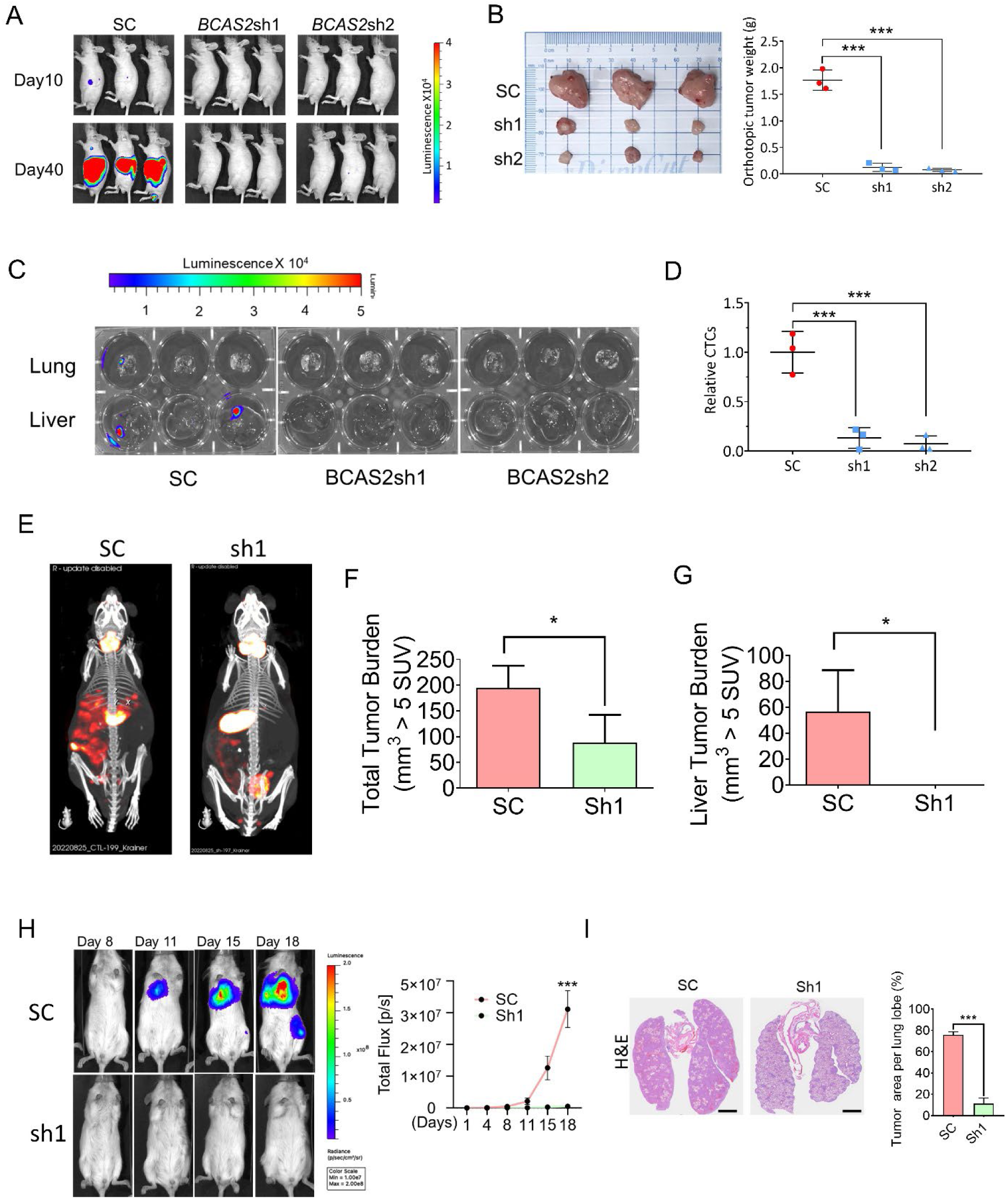
*BCAS2* knockdown suppresses PDAC tumor progression *in vivo*. (A) Luciferase bioluminescence imaging of pancreas orthotopic transplantation of MiaPaCa-2 cells with or without *BCAS2* knockdown. n = 3. (B) Primary tumors (left) and tumor weights (right) from pancreas orthotopic transplantation of MiaPaCa-2 cells with or without *BCAS2* knockdown. One-way ANOVA with Tukey multiple comparisons test. ***p < 0.001; n = 3. Error bars represent mean ± SD. (C) Luciferase bioluminescence imaging of lung tissues from pancreas orthotopic transplantation of MiaPaCa-2 cells with or without *BCAS2* knockdown. (D) Circulating tumor cells (CTC) quantified by qRT-PCR targeting human Alu-sequence and normalized by mouse *Actin* in the blood samples obtained from pancreas orthotopic transplantation of MiaPaCa-2 cells with or without *BCAS2* knockdown. One-way ANOVA with Tukey multiple comparisons test. ***p < 0.001; n=3. Error bars represent mean ± SD. (E) Serial SPECT/CT imaging of pancreas orthotopic transplantation of MiaPaCa-2 cells expressing sodium iodide symporter, with or without *BCAS2* knockdown. SC = 3, sh1 = 5. Maximum-intensity projection SPECT images were rendered from a scale of 0 – 20 SUV. (F and G) Total tumor burden (F) and liver tumor burden (G) quantified by volume above a threshold standardized uptake value (SUV) of 5 at week five following pancreas orthotopic transplantation, from (E). Unpaired two-tailed t-test, *p < 0.05, **p < 0.01; SC = 3, sh1 = 4. (H) Luciferase bioluminescence imaging (left) and quantification (right) of pulmonary metastatic foci following tail-vein injection of MiaPaCa-2 cells with or without *BCAS2* knockdown. Unpaired two-tailed t-test, ***p < 0.001; n = 10. (I) H&E staining and quantification of pulmonary metastatic foci from (H). Scale bar, 5 mm. Unpaired two-tailed t-test, ***p < 0.001; n = 10. Error bars represent mean ± SD.

We further employed single photon emission computed tomography/x-ray computed tomography (SPECT/CT) imaging, which enables precise visualization of tumor localization, metastatic lesions, and overall organ involvement.^36–38^ Consistent with our bioluminescence findings, orthotopic transplantation of MiaPaCa-2 cells stably expressing the sodium iodide symporter (NIS) SPECT reporter gene revealed a marked reduction in total tumor burden, as well as abdominal and hepatic metastases, in *BCAS2*-knockdown mice (Fig. 3E-3G).

Finally, to evaluate the role of BCAS2 in tumor metastasis independently of differences in primary tumor growth, we used an intravenous injection model of lung metastasis. *BCAS2* knockdown dramatically suppressed lung metastasis (Fig. 3H). Consistently, H&E staining showed significantly fewer and smaller metastatic lesions in *BCAS2* knockdown mice (Fig. 3I), which was supported by bioluminescence imaging analysis of lung tissues (Fig. S2). These results indicate that BCAS2 promotes PDAC tumor growth and metastatic progression *in vivo*.

### BCAS2 regulates specific alternative splicing events

We next examined splicing alterations upon *BCAS2* knockdown. RNA-seq analysis revealed that *BCAS2* knockdown predominantly affected cassette exon (CE) splicing events, which accounted for approximately 51% of all detected changes (Fig. 4A). Notably, *BCAS2* knockdown induced both increased and decreased exon inclusion, rather than causing a global impairment of spliceosome function (Fig. 4B and 4C).

**Figure 4.**
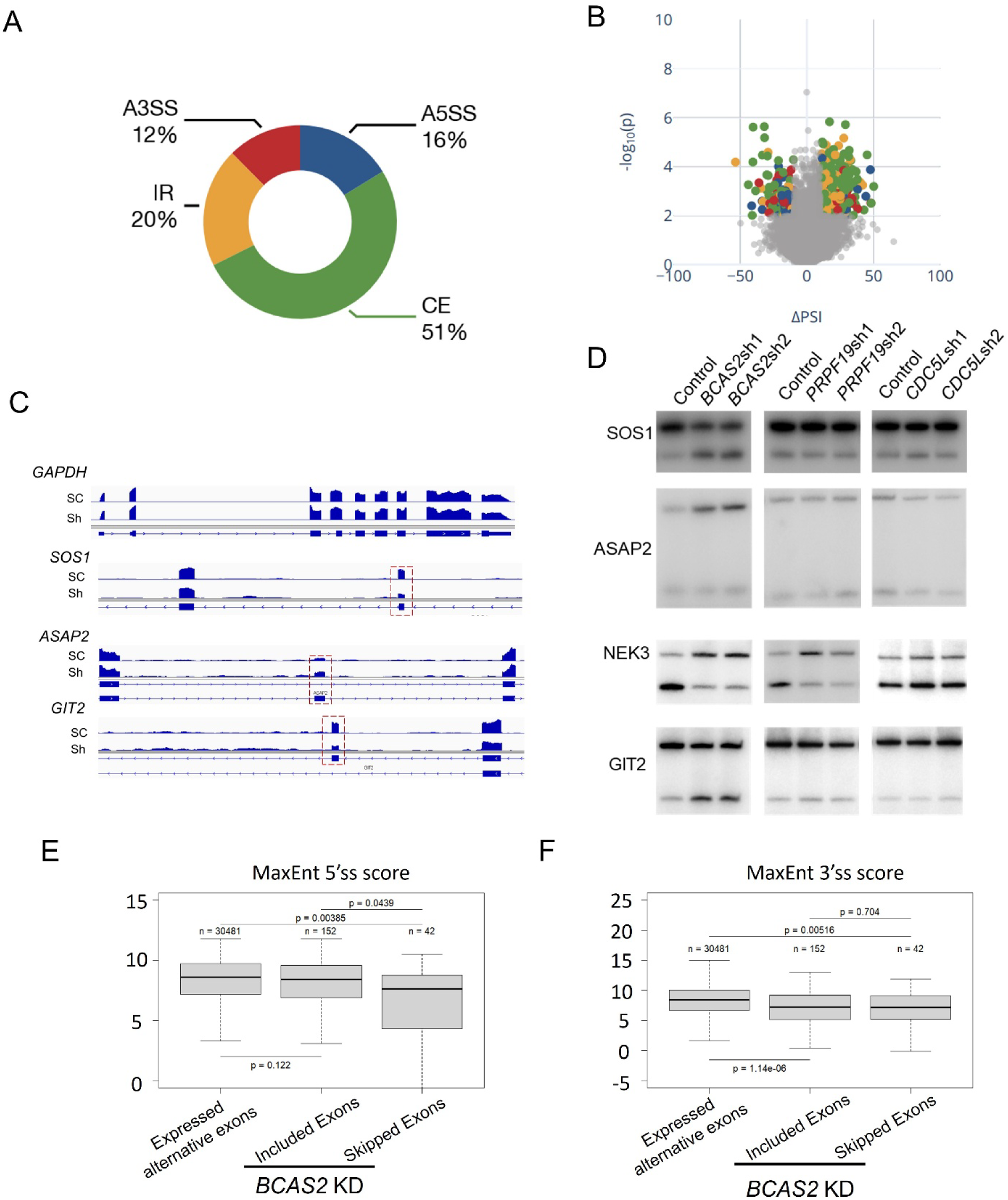
*BCAS2* knockdown causes specific alternative splicing changes. (A) Percentage of each alternative splicing event type altered in MiaPaCa-2 cells with or without *BCAS2* knockdown. (B) Volcano plots of splicing changes. PSI, percent spliced in. (C) Integrative Genomics Viewer (IGV) tracks showing alignment of RNA-seq reads of *GAPDH*, *SOS1*, *ASAP2*, and *GIT2* from RNA-seq of MiaPaCa-2 cells with or without *BCAS2* knockdown. Alternative exons are highlighted with red dashed boxes. (D) Radioactive RT–PCR analysis of splicing events in *SOS1, ASAP2, NEK3,* and *GIT2* in MiaPaCa-2 cells, following knockdown of *BCAS2, PRPF19,* and *CDC5L,* respectively. (E, F) Median 5′ (E) and 3′ (F) splice-site scores, calculated using MaxEnt for nucleotide sequences spanning the exon–intron boundaries, for expressed alternative exons and exons with increased or decreased inclusion, following *BCAS2* knockdown.

We validated a subset of *BCAS2*-regulated splicing events using radioactive RT-PCR (Fig. 4D and S3). As the NTC complex components are present stoichiometrically, we examined whether other core subunits influence the same BCAS2-dependent splicing events. Knockdown of *PRPF19* or *CDC5L* resulted in only minor alterations in BCAS2-regulated splicing events (Fig. 4D and S4). This disparity may reflect differential contributions of individual NTC components to the dynamic spliceosome conformational transitions.

We evaluated the splice-site strengths of BCAS2-regulated CEs. Compared with all expressed alternative exons, exons skipped upon *BCAS2* knockdown exhibited significantly weaker 5′ splice sites, whereas exons with increased inclusion showed comparable 5′ splice-site strength (Fig. 4E). Additionally, skipped exons exhibited significantly weaker 5′ splice sites than exons with increased inclusion. In contrast, 3′ splice-site strength did not differ between exons with increased and decreased inclusion (Fig. 4F), suggesting that 5′ splice-site strength is more closely associated with BCAS2-dependent exon regulation.

### BCAS2 regulated alternative splicing associated with Rho signaling

Pathway enrichment analysis of BCAS2-regulated alternative splicing events revealed significant enrichment of the RHO GTPase signaling pathway (Fig. 5A). Among these events, we identified an alternative splicing event in SOS1, a guanine nucleotide exchange factor (GEF).^39–41^ SOS1 is a RAS GEF that promotes GDP–GTP exchange on RAS family proteins, thereby activating MAPK/ERK and PI3K–AKT signaling pathways involved in cell proliferation and survival.^42, 43^ In addition, SOS1 regulates RHO-family GTPase signaling through its C-terminal Dbl homology (DH) and pleckstrin homology (PH) domains, which mediate RAC1 activation and control cytoskeletal remodeling, cell adhesion, and migration.^39^

**Figure 5.**
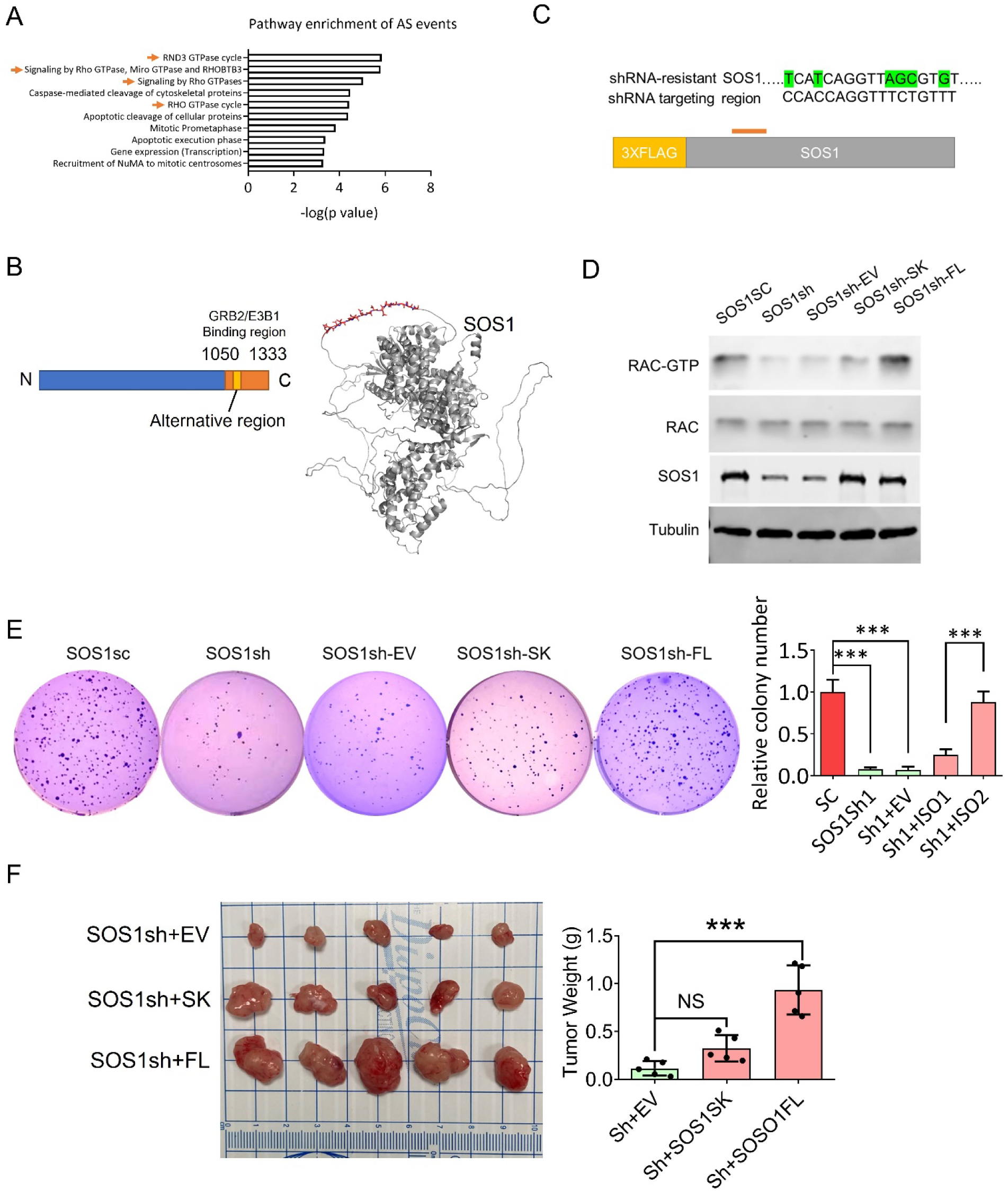
BCAS2 promotes Rho signaling by regulating *SOS1* alternative splicing changes. (A) Pathway-enrichment analysis of splicing changes from MiaPaCa-2 cells with *BCAS2* knockdown. (B) Schematic of *SOS1* full-length (FL) isoform (left) and AlphaFold-predicted SOS1-FL protein structure, with the alternative-exon peptide highlighted in red (right). (C) Construction of shRNA-resistant plasmids expressing SOS1 isoforms. (D) Western blot analysis of GTP-bound RAC, total RAC, SOS1, and Tubulin from MiaPaCa-2 cells with *SOS1* knockdown, followed by expression of individual SOS1 isoform cDNAs. (E) Soft-agar colony-formation assay (left) and quantification (right) of MiaPaCa-2 cells with *SOS1* knockdown, followed by expression of individual SOS1 isoform cDNAs. One-way ANOVA with Tukey multiple comparisons test. ***p < 0.001; n = 3. Error bars represent mean ± SD. (F) Tumors (left) and quantification of tumor weights (right) from subcutaneous xenografts generated by MiaPaCa-2 cells with *SOS1* knockdown, followed by expression of individual SOS1 isoforms. One-way ANOVA with Tukey multiple comparisons test. **p < 0.01; n = 3. NS, non-significant. Error bars represent mean ± SD.

We found that *BCAS2* knockdown suppressed inclusion of *SOS1* exon 21, which encodes a short peptide located in the C-terminal region (Fig. 4D and 5B). AlphaFold^44^ structural prediction indicates that this peptide lies within a flexible, unstructured region. We knocked down *SOS1* expression in MiaPaCa-2 cells and subsequently overexpressed shRNA-resistant *SOS1* isoforms to assess functional differences between them (Fig. 5C). *SOS1* knockdown significantly decreased active RAS and RAC levels (Fig. 5D and S5). Whereas both *SOS1* isoforms—exon 21-included (FL) and exon 21-skipped (SK)—rescued active RAS levels, the FL isoform displayed a markedly greater ability to increase RAC activation, consistent with previous studies demonstrating an essential role for the SOS1 C-terminus in RAC activation.^39^ Accordingly, MiaPaCa-2 cells expressing the *SOS1* FL isoform exhibited higher colony formation ability and tumor growth rate (Fig. 5E and 5F).

### Weak 5′ splice-site strength confers BCAS2-dependent regulation of *SOS1* exon 21

We next sought to determine how BCAS2 regulates *SOS1* exon 21 alternative splicing. Exon 21 has a relatively strong 3′ splice site, but a weak 5′ splice site (Fig. 6A). A *SOS1* minigene reporter comprising exons 20–22 and the flanking 150-bp intronic regions successfully recapitulated BCAS2-dependent regulation of exon 21 splicing (Fig. 6A). Strengthening the 5′ splice site abolished *BCAS2* knockdown-mediated exon 21 skipping, whereas weakening the 3′ splice site failed to restore BCAS2 regulation, suggesting that the weak 5′ splice site is critical for BCAS2-dependent regulation (Fig. 6A). To determine whether this regulation is sequence-dependent, we replaced the endogenous 5′ splice site with a distinct weak 5′ splice site, which retained *BCAS2* knockdown-induced exon skipping. To investigate whether BCAS2-mediated splicing regulation is context-dependent, we replaced the strong 5′ splice site of *SMN2* exon 7 with the weak *SOS1* exon 21 5′ splice site in an *SMN2* minigene reporter^45–47^ (Fig. 6B). Introduction of the weak *SOS1* 5′ splice site sensitized the reporter to *BCAS2* knockdown-induced exon skipping. These results identify weak 5′ splice-site strength as the primary determinant of BCAS2-dependent regulation of *SOS1* exon 21 splicing.

**Figure 6.**
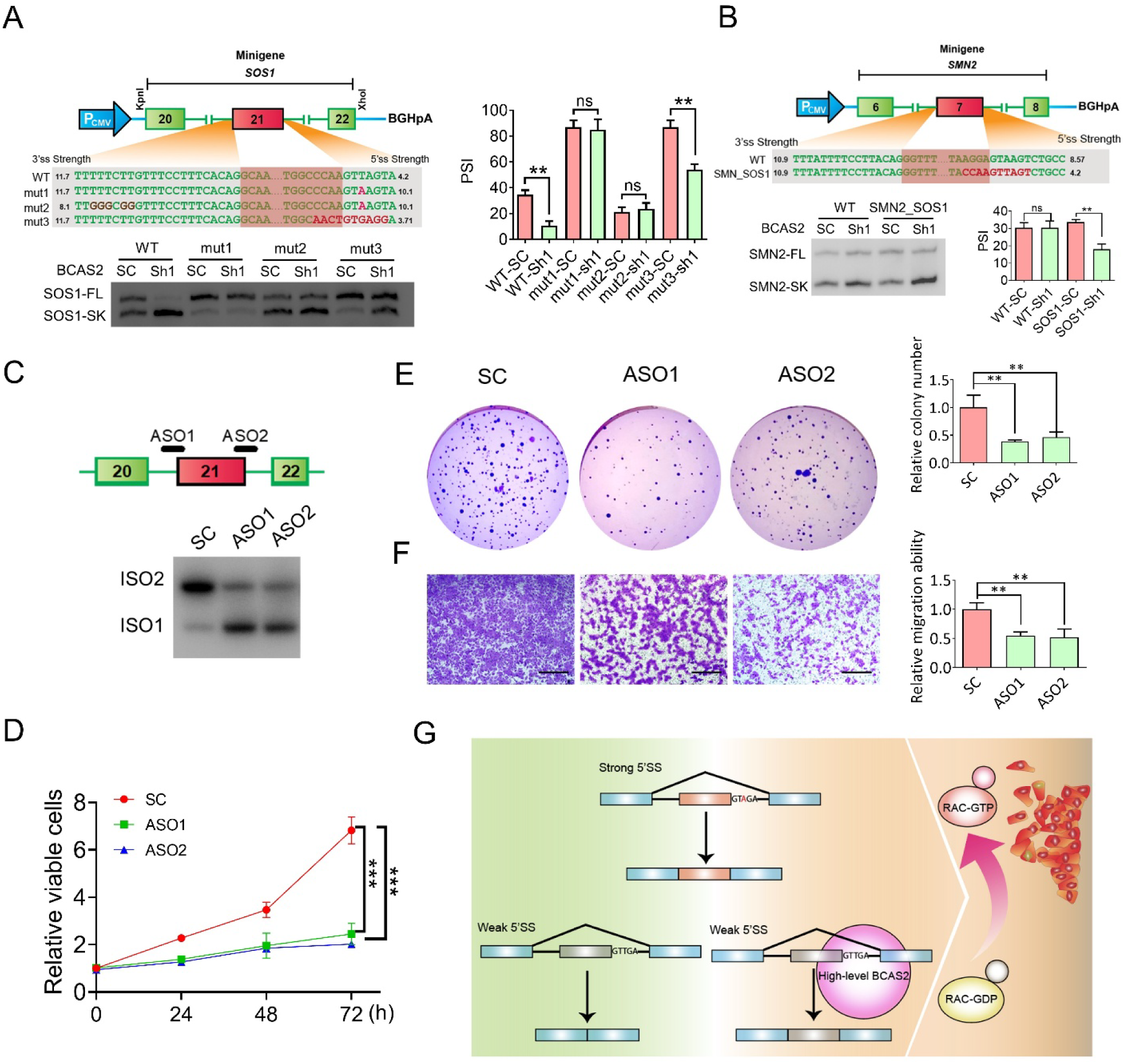
BCAS2 promotes *SOS1* exon 21 inclusion through weak 5′ splice site– dependent regulation. (A) Diagram and radioactive RT-PCR assays of *SOS1* minigenes in MiaPaCa-2 cells with or without *BCAS2* knockdown (left). The 5′ and 3′ splice sites were mutated to generate variants with different strengths. The percentage of exon 21 inclusion was quantified by radioactive RT–PCR (right). Unpaired two-tailed t-test. \*\**p* <0.01; n =3. Error bars represent mean ± SD. (B) Diagram of *SMN2* minigene assays in MiaPaCa-2 cells with or without *BCAS2* knockdown. The endogenous 5′ splice site of *SMN2* exon 7 was replaced with the weak 5′ splice site from *SOS1* exon 21. The percentage of exon 7 inclusion was quantified by radioactive RT–PCR. Unpaired two-tailed t-test. \*\**p* <0.01; n =3. Error bars represent mean ± SD. (C) Scheme of ASOs that modulate *SOS1* exon 21 inclusion (top), and radioactive RT–PCR of *SOS1* alternative splicing from MiaPaCa-2 cells transfected with ASOs. SC, scramble control. (D) Cell Counting Kit-8 (CCK-8) assays in MiaPaCa-2 cells transfected with ASOs. Unpaired two-tailed t test. ***p < 0.001; n=3. Error bars represent mean ± SD. (E) Soft agar colony formation assay (left) and quantification (right) of MiaPaCa-2 cells transfected with ASOs. One-way ANOVA with Tukey multiple comparisons test, **p < 0.01; n=3. Error bars represent mean ± SD. (F) Transwell assay (left) and quantification (right) of the cell migration properties of MiaPaCa-2 cells transfected with ASOs. One-way ANOVA with Tukey multiple comparisons test, **p < 0.01; n=3. Error bars represent mean ± SD. Scale bars, 200 μm. (G) Model illustrating BCAS2-mediated regulation of weak 5′ splice site–dependent alternative splicing in PDAC. Exons with strong 5′ splice sites undergo efficient splicing independently of BCAS2, whereas weak 5′ splice site-containing exons, such as *SOS1* exon 21, require BCAS2-dependent spliceosome regulation for efficient inclusion, which promotes RAC activation and contributes to PDAC progression.

Finally, we sought to modulate *SOS1* splicing with splice-switching antisense oligonucleotides (ASOs). ASOs targeting the exon 21 splice sites markedly suppressed exon 21 inclusion (Fig. 6C). Additionally, ASOs significantly suppressed SUIT-2 cell proliferation, colony formation, and migration (Fig. 6D-F). These results suggest that selective modulation of *SOS1* splicing provides a potential strategy to suppress PDAC progression.

## DISCUSSION

Dysregulated RNA splicing has emerged as a critical mechanism contributing to cancer progression.^5–9^ Recent studies have focused on mutations in spliceosomal components or altered expression of sequence-specific RNA-binding proteins.^5, 12–16, 19, 20^ How cancer cells exploit non-RNA binding spliceosome-scaffold components to regulate oncogenic splicing remains poorly understood. In this study, we identify BCAS2—a conserved component of the NTC complex—as a critical driver of pancreatic cancer progression. BCAS2 is upregulated in human PDAC and associated with poor clinical outcomes. Functional studies revealed that BCAS2 promotes PDAC cell proliferation, migration, tumor growth, and metastatic progression. Mechanistically, we showed that BCAS2 selectively regulates alternative splicing events enriched for weak splice-site features, including *SOS1* exon 21 inclusion, thereby activating RAC/RHO signaling.

Although BCAS2 is an essential NTC component required for spliceosome dynamics, we show that partial knockdown of *BCAS2* resulted in selective and bidirectional splicing changes, rather than widespread loss of splicing. BCAS2 preferentially regulates alternative exons characterized by suboptimal splice-site strength. Consistent with a previous study in HeLa cells^48^, we observed preferential splicing changes in cassette exons in *BCAS2*-knockdown PDAC cells.

Notably, exons with decreased inclusion following *BCAS2* knockdown exhibited significantly weaker 5′ splice sites than either exons with increased inclusion or expressed alternative exons. In contrast, 3′ splice-site strength did not differ significantly between the increased- and decreased-inclusion groups, revealing a preferential association between weak 5′ splice sites and BCAS2-dependent exon inclusion. We propose that BCAS2 facilitates spliceosome activation on kinetically disadvantaged spliceosomal complexes assembled on exons with suboptimal splice sites. According to this model, weak 5′ splice sites reduce the efficiency or stability of early spliceosome assembly, rendering subsequent conformational transitions more dependent on BCAS2. In contrast, strong 5′ splice sites efficiently support spliceosome assembly and catalytic activation, with reduced dependence on BCAS2.

Notably, the failure of *PRPF19* or *CDC5L* knockdown to recapitulate BCAS2-dependent splicing changes suggests that individual NTC components make distinct contributions to spliceosome conformational transitions, rather than functioning solely as static structural components. This functional specificity may reflect differences in their positions within the spliceosome, interaction partners, or roles during spliceosome remodeling. Another possibility is that BCAS2 has regulatory effects on splicing as a separate physical entity from the NTC complex. Future structural and biochemical studies will be required to define how BCAS2 facilitates spliceosome conformational transitions on weak splice-site substrates.

Using *SOS1* exon 21 as a model, we demonstrated that strengthening the 5′ splice site eliminates regulation by BCAS2, whereas replacing it with a different weak 5′ splice site retained sensitivity to *BCAS2* knockdown. Moreover, introducing the weak *SOS1* exon 21 5′ splice site into a heterologous exon context was sufficient to confer BCAS2-dependent regulation. These findings support a model in which BCAS2 facilitates the processing of alternative exons with intrinsically weak 5’ splice sites, rather than recognizing specific exon sequences.

Our study identified *SOS1* alternative splicing as an important downstream effector of BCAS2-mediated oncogenic splicing. Although both *SOS1* isoforms effectively activated RAS signaling, the exon-21-included isoform preferentially promoted RAC activation. These findings reveal a previously unrecognized layer of regulation controlling the balance between RAS- and RAC-dependent signaling pathways in PDAC.

Therapeutically modulating aberrant splicing is an emerging strategy for diseases driven by pathogenic RNA isoforms.^49–51^ While direct inhibition of core spliceosome components can be limited by toxicity, due to their essential cellular functions, targeting individual cancer-promoting splicing events may provide greater specificity. Our preclinical evaluation showed that ASO-mediated *SOS1* exon 21 skipping suppressed PDAC cell proliferation and migration. These findings provide proof-of-principle evidence that downstream splicing events controlled by a spliceosome-scaffold component can be therapeutically modulated.

In summary, our study identified an oncogenic function of the spliceosome-scaffold component BCAS2 in pancreatic cancer. BCAS2 selectively enhances weak 5’ splice site recognition to support tumor progression. The identification of the BCAS2–*SOS1* splicing–RAC signaling axis provides new mechanistic insight into how a dysregulated spliceosome structural component contributes to PDAC pathogenesis (Fig. 6G).

## METHOD

### Analysis of Public Datasets

mRNA expression levels of *BCAS2, PRPF19,* and *CDC5L*, along with relevant clinical information, were determined using the GEPIA Genome Atlas (TCGA) and Genotype-Tissue Expression (GTEx) databases. Expression of *BCAS2* across different tissues was determined by re-analysis of RNA-seq performed of tissue samples from 95 human individuals representing 27 different tissues.^52^

### Western Blotting

Cells were harvested and lysed in RIPA buffer (50 mM Tris-HCl, pH 7.4; 150 mM NaCl; 0.1% SDS; 1% NP-40; 0.5% sodium deoxycholate; 1 mM PMSF; and protease inhibitor cocktail [Roche]) by incubation on ice for 30 minutes. Western blotting was performed using the following primary antibodies at 1:1000 dilution: rabbit anti-BCAS2 (Proteintech, 10414-AP), rabbit anti-SOS1 (Cell Signaling Technology, 5890), rabbit anti-Tubulin (Proteintech, A01203), rabbit anti-RAC (Cell Signaling Technology, 2465S). Blots were imaged and quantified using an Odyssey Infrared Imaging System (LI-COR).

### Detection of Active RAS and RAC Small-GTPases

Cells were lysed and processed using immunoprecipitation kits for detection of active RAS (Cell Signaling Technology, 8821) and RAC (Cell Signaling Technology, 8815) following the manufacturer’s instructions. Levels of active RAS and RAC small-GTPases were determined by western blotting after immunoprecipitation.

### Transwell Migration Assays

Cell migration was measured on Transwell chamber plates (24-well format; 8-μm pore size; Corning Costar, New York, USA). Images of cells on the lower face of the filter were captured in fields at 10× magnification. To quantify the cells that had migrated across the filters, we used ImageJ (National Institutes of Health, USA) to measure the cell area on the lower face of the filter.

### RT-PCR

Total RNA was isolated using Trizol (Invitrogen). Then RT-PCR was performed using PrimeScript™ RT Reagent Kit (Takara, CA, USA) according to the manufacturer’s manual. Radioactive PCR was conducted using ^32^P-α-dCTP, 1.25 U of AmpliTaq (Invitrogen), and 26 cycles. Products were separated by 5% PAGE, and the bands were detected using a Typhoon FLA 7000 instrument (GE Healthcare, IL, USA). Quantitative RT-PCR was performed on a QuantStudio 6-flex Real-time PCR instrument (Applied Biosystems, MA, USA). Expression levels were normalized to *GAPDH*.

### Soft-Agar Colony Formation

Cells (10^3^ cells/well) were suspended in an upper layer of 0.3% agarose (ThermoFisher Scientific) in DMEM with 10% FBS. The plate was placed at room temperature until the top agarose solidified. The plate was then incubated at 37 °C/5% CO_2_ for at least 3 weeks, before staining with crystal violet. Visible colonies were then counted.

### Mouse Tail-Vein Assay

Cells stably expressing luciferase and transduced with either scramble control (SC) or *BCAS2* shRNA were injected into the tail vein of NOD scid gamma (NSG) mice. Bioluminescence imaging was initiated 5 d after injection. Mice were injected with luciferin (300 mg/kg) 5 min before imaging, anesthetized with 3% isoflurane, and imaged using an IVIS Spectrum Imaging System (Caliper).

### Transplantation Models

For subcutaneous model, cells were suspended in 100 μl PBS and injected into each flank of BALB/c nu/nu nude mice (male, 6 weeks of age). Tumor growth was measured at 3-day intervals after injection using a caliper, and tumor volume was calculated according to the following formula: length × width^2^ × 0.5. For orthotopic engraftment, cells were harvested and washed with ice-cold PBS, and resuspended in 50 μL of Matrigel diluted 1:1 with cold PBS. Cells were injected into the tail of the pancreas using a 29-gauge insulin syringe. Successful injection was confirmed by the formation of a fluid bleb without evidence of intraperitoneal leakage. Harvested tumor tissues were fixed in 10% buffered formalin, embedded in paraffin, sectioned, and stained.

### SPECT imaging

*In vivo* SPECT images were acquired with a Mediso nanoScan SPECT/CT scanner (Mediso USA, Arlington, VA, USA). Mice were injected intravenously via the tail vein with a nominal activity of 50 MBq [^99m^Tc]sodium-pertechnetate. After conscious uptake, mice were anesthetized and a CT scan (360 projections at 50 kVp and 111 µAs exposure) acquired for anatomical reference and attenuation correction. A whole-body pinhole SPECT scan was then acquired 60 min after radiotracer injection. CT images were reconstructed using a filtered back-projection algorithm with a cosine filter to an isotropic voxel size of 250 µm. SPECT images were reconstructed using a 3D iterative algorithm optimized for high dynamic range, with 48 iterations and three subsets to a 128×128 matrix with a 260 µm isotropic voxel size. Attenuation, decay and scatter corrections were applied. SPECT/CT images were processed and analyzed with VivoQuant 4.0 software (inviCRO, Boston, MA, USA) as described previously^36^. Briefly, SPECT images were converted to units of SUV, and an arbitrary threshold of 5 SUV was applied to highlight regions of increased uptake. Referencing the CT image, stomach and other sites of endogenous pertechnetate uptake were manually removed from the threshold region, and the remaining total viable tumor volume was quantified. Additionally, viable tumor volume located only within the liver was quantified.

### Immunohistochemistry (IHC)

Human Tissue microarrays were obtained from US Biomax (PA483e, PA1002b, BIC14011b). Mouse tissue samples were fixed in 4% PFA/PBS (EMS, Hatfield, PA, USA) and embedded in paraffin. Six-µm sections were deparaffinized and rehydrated. Endogenous peroxidase was blocked in H_2_O_2_/methanol for 15 min. Slides were then boiled in 10 mM sodium citrate, 0.05% Tween 20 for 5 min under pressure (for protein antigens). Slides were blocked for 30 min with peptide blocking solution (Innovex Biosciences, Richmond, CA, USA) and incubated with the indicated primary antibodies overnight at 4 °C. For IHC analysis, the signal was visualized with horseradish peroxidase (HRP)-labeled anti-rabbit polymer and DAB (DAKO, Carpinteria, CA, USA). Slides were counterstained with hematoxylin (Sigma-Aldrich) and mounted with Limonene mounting medium (Abcam, Cambridge, UK).

### Antisense Oligonucleotide Transfections

2’-methoxyethyl (2’-MOE)-modified phosphorothioate ASOs were purchased from Integrated DNA Technologies (IDT, IA, USA). ASOs were delivered to cells using Lipofectamine 2000 (Invitrogen, 11668019) at a final concentration of 60 nmol/L. Cells were harvested after 72 h for RT-PCR and western blotting analysis.

### Gene Expression Analysis

RNA was extracted using Trizol as described above. RNA libraries were generated using NEBNext Ultra II Directional RNA Library Prep Kit for Illumina (NEB), following the manufacturer’s instructions. The libraries were sequenced on an Illumina NextSeq 2000 (150 bp, paired-end). Raw Illumina sequencing reads were aligned to genome assembly GRCh38 using STAR 2.5.3a. ΔPSI and FDR values of splicing changes were produced by using PSI-Sigma v1.9j.^53^ The Database for Annotation, Visualization and Integrated Discovery (DAVID) v6.8 was used to identify enriched Kyoto Encyclopedia of Genes and Genomes (KEGG) pathways for up- and down-regulated genes, respectively.^54^

Splice-site strength was calculated using the MaxEnt method^55^ for nucleotide sequences spanning the 5′ and 3′ exon–intron boundaries, extracted from the GRCh38 primary-assembly genome sequence and retaining only canonical (GT–AG or GC–AG) splice sites. Alternative exons were identified computationally from the Ensembl GRCh38 gene annotation^56^ as internal exons of a GENCODE/Ensembl “basic” transcript that were either skipped or replaced by a different exon between the same flanking splice sites in at least one other annotated transcript of the same gene, irrespective of whether that other transcript was itself annotated as “basic”. BCAS2-regulated exons were identified by PSI-Sigma^57^ based on changes in exon inclusion following BCAS2 knockdown. Expressed alternative exons were defined as Ensembl-annotated alternative exons for which both introns flanking the exon were supported by more than 10 uniquely mapping reads in MiaPaCa-2 RNA-seq data (STAR SJ.out.tab^58^). Splice-site scores were compared using the Wilcoxon rank-sum test.

## Acknowledgments

We gratefully acknowledge the assistance of the Cold Spring Harbor Laboratory (CSHL) Animal, Animal and Tissue Imaging, Microscopy, Flow Cytometry, Antibody, Next-Generation Sequencing, and Histology Shared Resources. This work was supported by National Cancer Institute (NCI) Program Project Grant P01CA13106 Project 2 (to A.R.K.); the Carol M. Baldwin Foundation and start-up funds from the Renaissance School of Medicine at Stony Brook University (to L.W.); and in part by American Cancer Society Institutional Research Grant 21-143-01-IRG. CSHL Shared Resources used in this study were supported in part by NCI Cancer Center Support Grant P30CA045508.

## Author Contributions

Conceptualization: L.W., A.R.K.; Methodology: L.W., M.W., Y. I., A.J.K., K.-T. L., J.R.M., S.K.L., A.R.K.; Investigation: L.W., M.W., Y. I., A.J.K., K.-T. L., J.R.M.; Writing: L.W., M.W., Y. I., J.R.M., S.K.L., A.R.K.; Resources: L.W., A.R.K.; Supervision: L.W., A.R.K.

---

**Figure S1.**
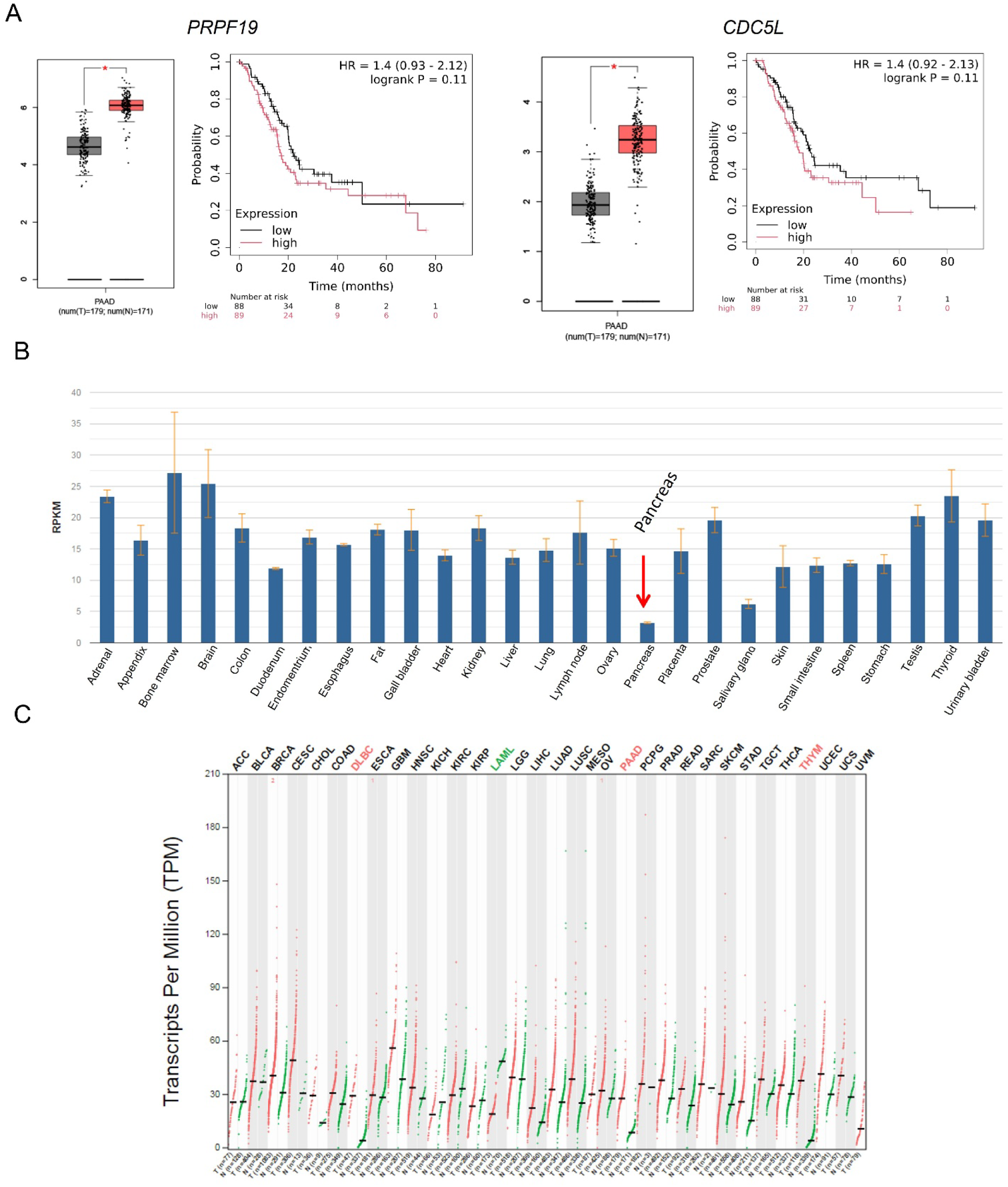
Elevated expression of NineTeen Complex core components in PDAC. (A) *PRPF19* and *CDC5L* expression in normal pancreatic samples and pancreatic cancer from TCGA RNA-sequencing data, and Kaplan–Meier survival analysis of PDAC patients with different expression levels. (B) *BCAS2* expression from 27 different tissues of 95 human individuals. (C) *BCAS2* expression across human cancers and normal tissues from TCGA and GTEx datasets. Expression is shown as TPM, with tumor samples in red and normal samples in green. Differential expression was determined by ANOVA with Benjamini–Hochberg FDR correction. Significantly upregulated (red) and downregulated (green) cancer types are highlighted.

**Figure S2.**
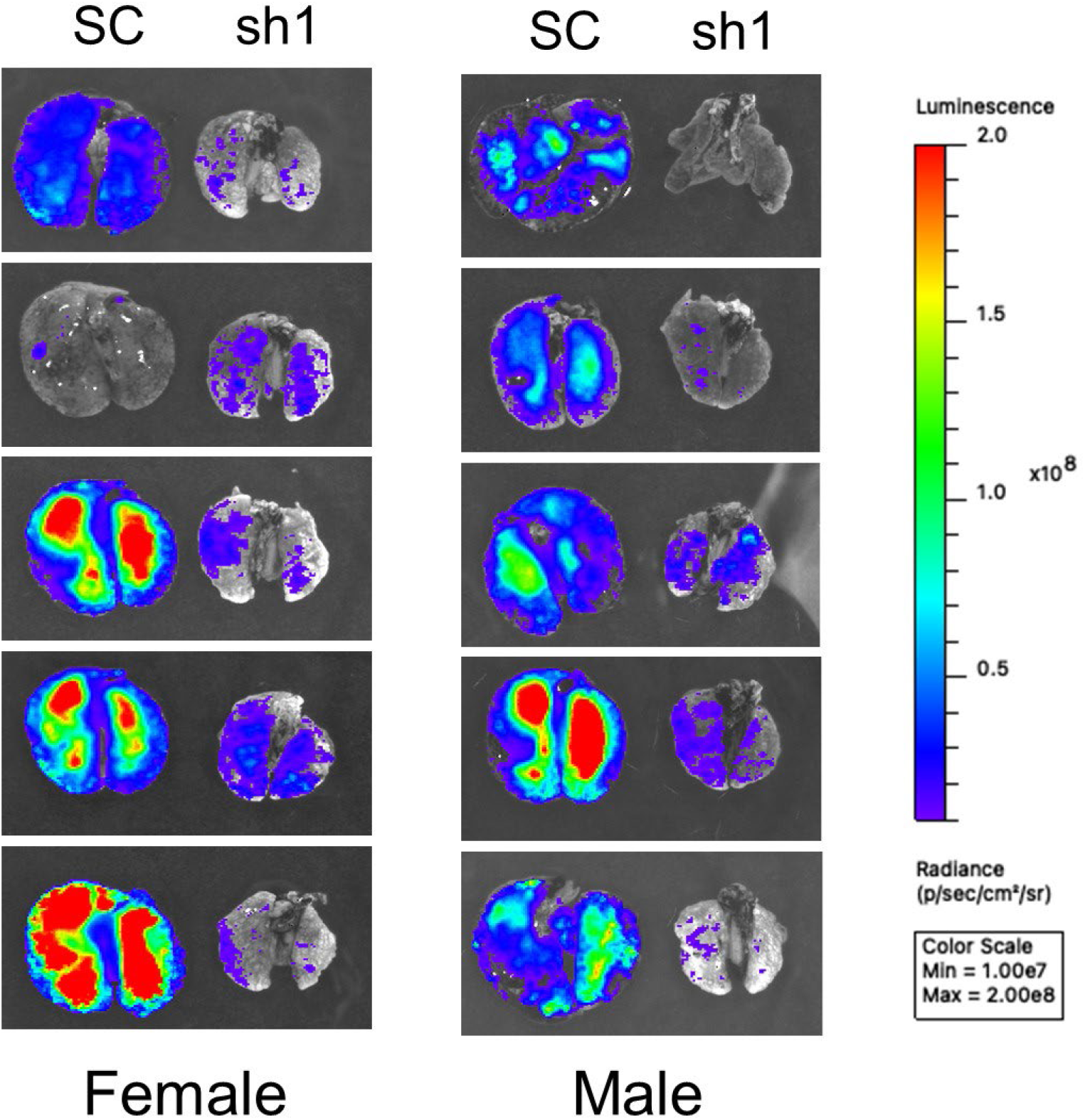
*BCAS2* knockdown suppresses PDAC tumor lung metastasis. Luciferase bioluminescence imaging of lung tissue from mice with tail-vein injection of MiaPaCa-2 cells with or without *BCAS2* knockdown.

**Figure S3.**
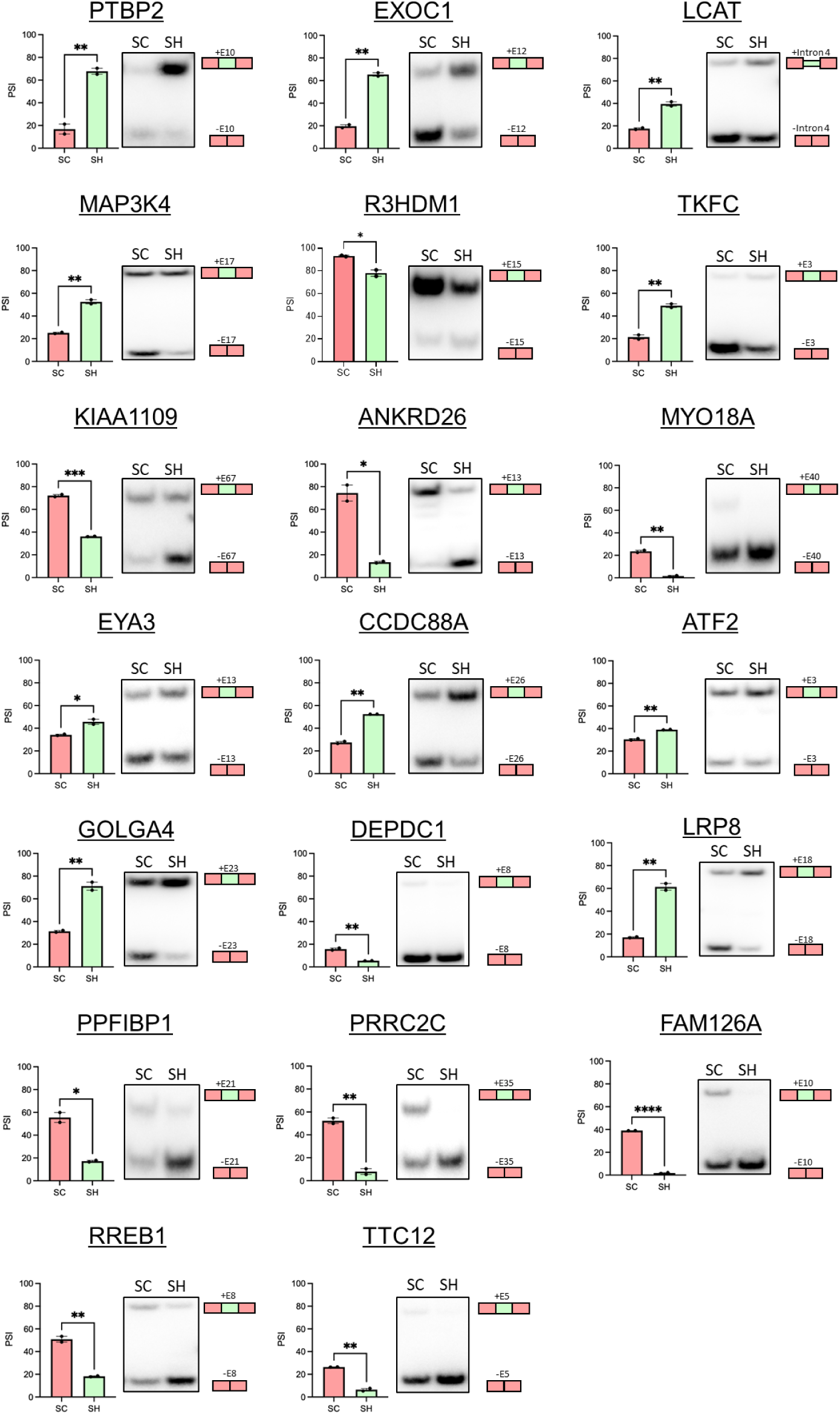
*BCAS2* knockdown regulates alternative splicing. Quantification (left) and radioactive RT-PCR (right) results of alternative splicing in MiaPaCa-2 cells with or without *BCAS2* knockdown. Unpaired two-tailed t-test. \**p* <0.05, \*\**p* <0.01, \*\*\**p* <0.001. Error bars represent mean ± SD.

**Figure S4.**
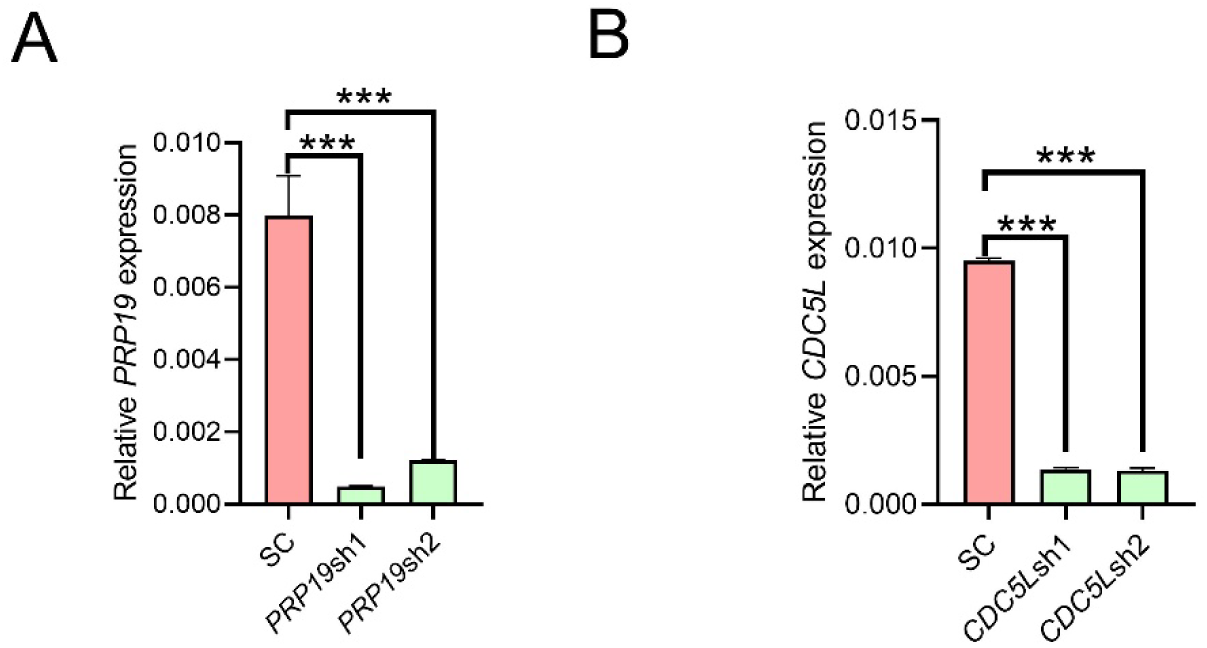
*PRP19* and *CDC5L* knockdown in MiaPaCa-2 cells. (A) Real-time qRT-PCR quantification of *PRP19* and expression in MiaPaCa-2 cells with or without *PRP19* knockdown. (B) Real-time qRT-PCR quantification of *CDC5L* expression in MiaPaCa-2 cells with or without *CDC5L* knockdown. One-way ANOVA with Tukey multiple comparisons test. ***p < 0.001; n=3. Error bars represent mean ± SD.

**Figure S5.**
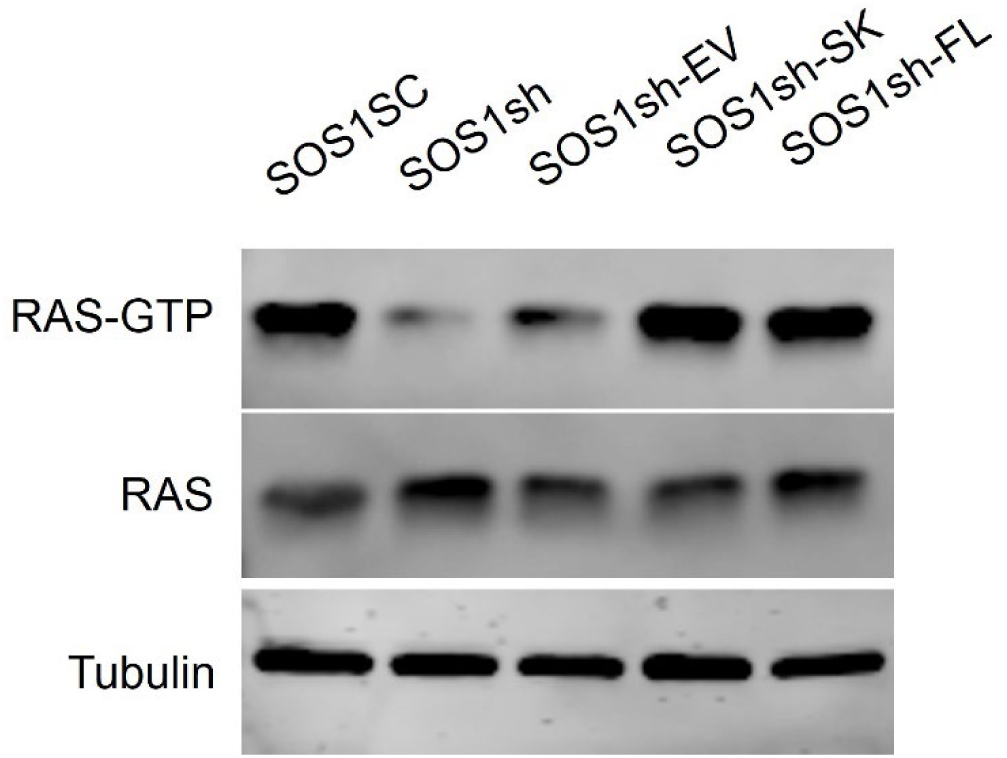
*SOS1* isoforms similarly restore RAS activation following *SOS1* knockdown. Western blot analysis of GTP-bound RAS, total RAS, and Tubulin from MiaPaCa-2 cells with *SOS1* knockdown, followed by expression of individual *SOS1* isoforms.

